# Optogenetic RNA relocalization reveals causal neural function *in vivo*

**DOI:** 10.64898/2026.11.17.767469

**Authors:** Sa Cai, Boxiong Deng, Maylin L. Fu, Alisilani Alusa, Praveen Prakhar, Gong-Her Wu, Alexander A. Choi, Dongsheng Han, Yinglin Situ, Mengting Han, Yanyu Zhu, Xiang Wu, Wah Chiu, Guosong Hong, Sui Wang, Lei S. Qi

## Abstract

RNA localization dictates where translation occurs, yet its disease role is unclear because most assays manipulate abundance, not position. We develop an optogenetic CRISPR-dCas13 system (Opto-CRISPR-TO) that repositions endogenous RNAs with subcellular precision, second-scale kinetics, and reversibility, enabling causal tests *in vivo*. In cells, Opto-CRISPR-TO routes diverse RNAs to organelles and ribonucleoprotein granules with multiplex, bidirectional control, revealing distinct RNA behaviors in centrosomes and P-bodies. In mouse retina, acute (4-hour) mislocalization of *Actb* mRNA redirects β-actin protein, remodels synaptic architecture and cell positioning, and persistently impairs vision. Thus, RNA location governs proteome topology and neural circuit output, and brief mislocalization drives lasting dysfunction. The approach generalizes across targets, compartments, and cell types, establishing a framework for probing RNA localization mechanisms in development, plasticity, and degeneration.

## INTRODUCTION

Spatial RNA organization is a fundamental layer of gene regulation that shapes where and when proteins are made (*1–5*). Although thousands of transcripts exhibit subcellular patterning across organisms, the functional and pathological consequences of RNA mislocalization remain poorly understood (*6–8*). A prevailing view holds that RNA abundance dominates phenotype, relegating localization to a secondary role (*9*). Yet abundance is experienced locally: the same number of transcripts at the wrong place can rewire translation, signaling, and organelle homeostasis (*10*, *11*). Indeed, mounting evidence links aberrant RNA localization to physiological functions and diseases (*12*, *13*), including neurological dysfunction and degeneration (*14*, *15*). However, causal studies linking RNA organization to function *in vivo* have been scarce.

High-content profiling has revealed that many transcripts occupy stereotyped subcellular niches, linking RNA function to local translation and organelle homeostasis. However, these atlases remain largely correlational, and tools that enable on-demand movement of endogenous RNAs to test causality between localization and phenotype are needed. Traditional RNA perturbations rely on genetic rewiring, such as deleting or swapping 3’ untranslated region (UTR) regulatory elements of RNA localization or inserting MS2 repeats bound by the fusion MS2 coat protein (MCP) and subcellular compartment protein for RNA relocalization. These traditional methods are powerful but labor-intensive, permanently alter the RNA, and offer limited reversibility or subcellular addressability in intact tissues (*11*,*16–18*). We recently introduced CRISPR-mediated transcriptome organization (CRISPR-TO), which repositions endogenous transcripts without changing their sequence via chemical-inducible dimerization to recruit dCas13 complexes (*19*). While effective in cells, drug induction imposes slow on/off kinetics, affords little spatial resolution, and faces potential delivery/penetration constraints that hinder *in vivo* tests of RNA localization-associated pathology.

Optogenetics modulates protein state with light, providing fast dynamics, full reversibility, subcellular resolution, and multiplexing via a broad palette of light-sensitive modules (*20*, *21*). These attributes are exactly what RNA-localization biology demands: second-to-minute control, patterned illumination, and noninvasive deployment in tissue. To date, however, optogenetics has only been applied to induce RNA clustering or condensation (*22*). Utilizing optogenetics for dynamic, programmable control of endogenous RNAs in distinct subcellular compartments with different cellular functions with high spatiotemporal precision is urgently needed.

Here we build an optogenetic CRISPR-TO (Opto-CRISPR-TO) that overcomes these technical limitations, enabling rapid, reversible, and spatially confined repositioning of endogenous RNAs in cultured cells and *in vivo*. This capability lets us move beyond atlases to causal examination, showing that acute, local misplacement of a single endogenous mRNA is sufficient to reprogram protein distribution, remodel circuitry, and impair function, thereby establishing local RNA localization-associated pathology as a first-order determinant of phenotype.

## RESULTS

### Opto-CRISPR-TO targets diverse endogenous RNAs across cell types

We developed an optogenetic CRISPR-based tool for RNA transport and organization to control the localization of endogenous RNAs. The system uses a catalytically inactive *Prevotella* sp. P5–125 Cas13 protein (dPspCas13b) fused to one half of a light-dimerizable module, which binds the target RNA recognized by a guide RNA (gRNA) (*23*). The other half of the module is fused to a subcellular anchor or a motor protein. Upon illumination, the two modules dimerize, recruiting the target RNA to the desired destination (**Figure 1A**).

**Figure 1.**
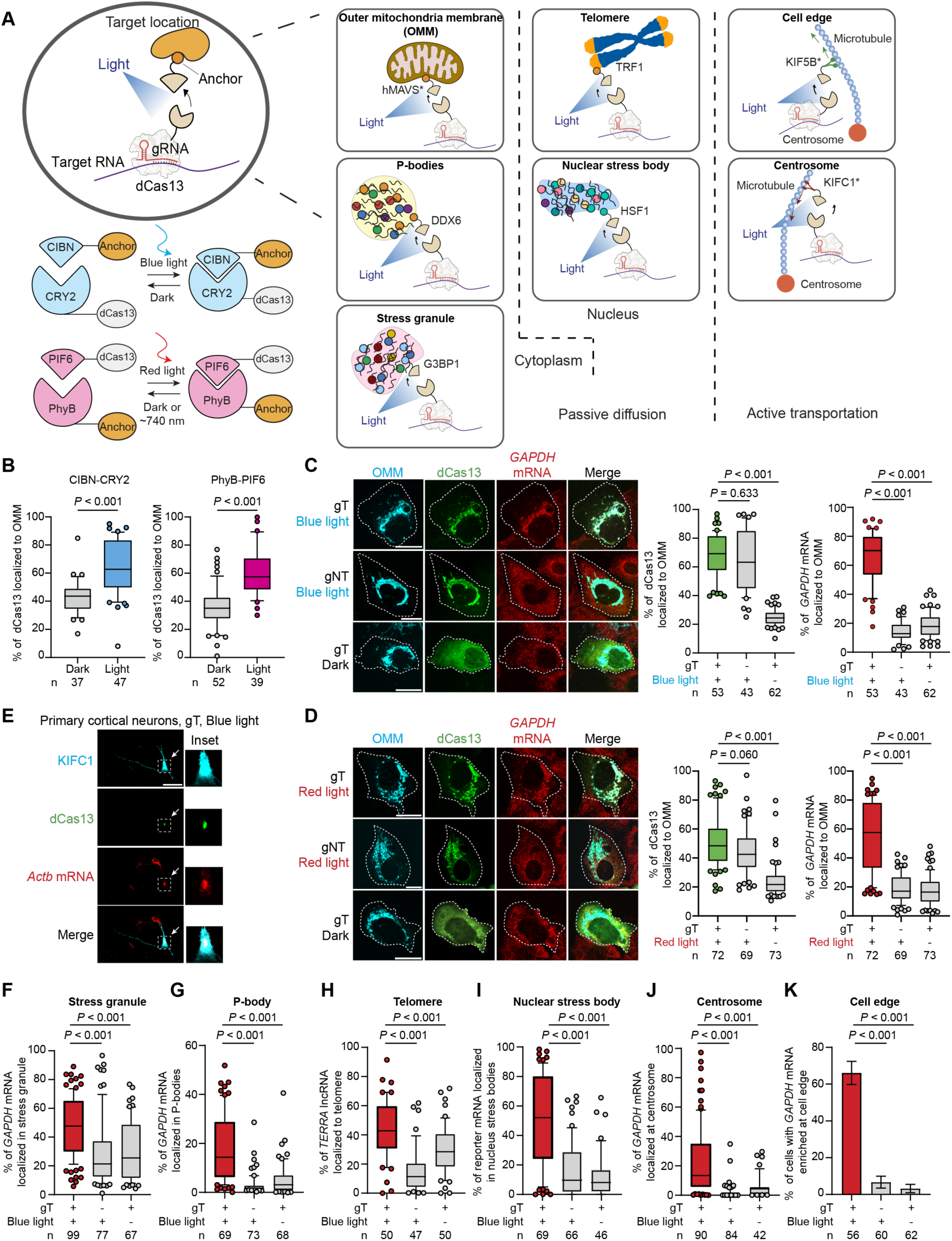
Opto-CRISPR-TO localizes different RNAs to varied subcellular compartments in multiple cell types. (**A**) Schematic of light-controlled RNA localization through tethering to subcellular anchors or active transport by motor proteins. (**B**) Percentage of dCas13 localized to the outer mitochondrial membrane (OMM) in individual HeLa cells under different light treatments for optogenetic dimerization proteins dCas13-CRY2/CIBN-hMAVS* and dCas13-PIF6/PhyB-hMAVS*. (**C** and **D**) Representative images and quantification of endogenous *GAPDH* mRNA and dCas13 protein localization to the OMM in HeLa cells using blue-light-triggered (**C**) or red-light-triggered (**D**) Opto-CRISPR-TO. White dotted lines indicate cell boundaries. Scale bars: 20 µm. (**E**) Representative images showing endogenous *Actb* mRNA recruitment to the centrosome (minus end of microtubules) in primary mouse cortical neurons following blue-light stimulation. Scale bar: 50 µm. (**F**–**J**) Quantification of blue-light-induced RNA localization: *GAPDH* mRNA to stress granules (**F**) and P-bodies (**G**), *TERRA* RNA to telomeres (**H**), *24×GCN4* reporter RNA to nuclear stress bodies (**I**), and *GAPDH* mRNA to the centrosome (**J**). Measurements were performed in individual cells (**F**, **G**, and **J**) or nuclei (**H** and **I**). (**K**) Percentage of transfected cells with *GAPDH* mRNA enriched at the cell periphery. Bars and error bars were presented as mean ± s.e.m. Localization percentages in (**B**–**D**) and (**F**–**J**) were calculated from fluorescence intensities. *P* values were calculated using two-tailed unpaired t-tests (**B**), one-way ANOVA followed by Tukey’s HSD post hoc tests (**C**, **D**, and **F**–**J**), and two-sided Fisher’s exact tests (**K**). Boxplots show the median, 25th–75th percentiles (boxes), 10th–90th percentiles (whiskers), and individual values beyond the whiskers. n indicates the number of cells or nuclei analyzed in each group. gT, targeting guide RNA; gNT, non-targeting guide RNA.

To build a robust and versatile system, we systematically evaluated four optogenetic pairs responsive to either blue or red/far-red light (CRY2-CIBN, PhyB-PIF6, DrBphP-Affibody, and BphP1-QPAS1) (*24–28*). We assessed their dimerization efficiency in HeLa cells by tethering one module to dCas13 and the other to an outer mitochondrial membrane (OMM) anchor (mitochondrial antiviral-signaling peptide, MAVS*) (*8*). Our screening revealed that the fusion orientation of the optogenetic domains was critical for function. For the blue-light-inducible CRY2-CIBN, fusing CRY2 to dCas13 yielded rapid and reversible recruitment to the OMM, whereas the reverse orientation was non-functional (**Figure 1B**). Similarly, for the red-light-inducible PhyB-PIF6, only the dCas13-PIF6 configuration mediated robust light-dependent dimerization that was reversible with far-red light (**Figure 1B**). Other tested pairs showed weak or no activity. We selected dCas13-CRY2/CIBN and dCas13-PIF6/PhyB for their efficient and specific light-dependent control, establishing a multiplex optogenetic platform for manipulating RNA localization.

We next validated that Opto-CRISPR-TO could relocalize endogenous RNAs using single-molecule RNA FISH (smiFISH) for visualization (*29*, *30*). Using blue-light CRY2-CIBN and gRNAs binding to the *GAPDH* 3’ UTR, *GAPDH* mRNA was enriched at the OMM to 65.9% versus 13.9% (non-targeting gRNA, gNT, light) and 18.8% (targeting gRNA, gT, dark), while dCas13 itself was recruited to the OMM in both illuminated groups (**Figure 1C**). With red-light PIF6–PhyB, *GAPDH* mRNA reached 55.1% at the OMM versus 19.0% (gNT, light) and 18.2% (gT, dark) in controls (**Figure 1D**).

By simply exchanging the gRNA, we next relocated other endogenous transcripts, including *Actb* and mRNAs encoding mitochondrial oxidative phosphorylation subunits (*SDHC*, *SDHD*, and *NDUFS2*), to the OMM (**Figure S1A-D**) (*31*). Furthermore, the system was effective across different cell lines, efficiently enriching *GAPDH* mRNA at the OMM in human embryonic kidney (HEK293T) cells and *Actb* mRNA in mouse neuroblastoma (N2A) cells (**Figure S1E-F**). Unless noted, subsequent experiments use the blue-light system.

We adapted the system for active, motor-driven transport of RNA in primary mouse cortical neurons. We targeted endogenous *Actb* mRNA and recruited it to a truncated minus-end-directed motor protein KIFC1* (*32*). Upon 4-hour blue-light stimulation, both dCas13 and its bound *Actb* mRNA were robustly transported to and enriched around the centrosome. This transport was specific, as it did not occur in control neurons that used gNT or were kept in the dark (**Figure 1E**).

### Opto-CRISPR-TO redirects endogenous RNAs to diverse cytoplasmic and nuclear compartments

We extended targeting beyond mitochondria by fusing the light dimerization partner to diverse compartment anchors or motor proteins (**Figure 1A**). In the cytoplasm, we targeted endogenous *GAPDH* mRNA to stress granules by anchoring to G3BP1 (*33*), observing robust enrichment in HeLa cells under sodium arsenite-induced stress (47.8% vs 29.1%/31.1% controls; **Figure 1F**; **S2A-B, J**). Similarly, anchoring to the P-body component protein DDX6 successfully sequestered *GAPDH* mRNA into P-bodies (17.4% vs 3.1%/4.9% controls; **Figure 1G**; **S2A**, **C**, **K**) (*34*).

For nuclear targeting, we determined that three nuclear localization signals (3xNLS) were required for efficient import of the dCas13-CRY2 fusion. Using this nuclear-localized system, we redirected the non-coding RNA *TERRA* to telomeres by anchoring to TRF1 (*35*). This resulted not only in significant *TERRA* enrichment but also in larger telomere puncta, supporting a positive role for *TERRA* in regulating telomere size (**Figure 1H**; **S2D-E**, **L**) (*36*). We also successfully localized a 24xGCN4 reporter transcript to nuclear stress bodies by anchoring to HSF1 (49.6% vs 16.8%/13.4%; **Figure 1I**; **S2D**, **F**, **M**) (*37*).

For active transport, the platform enables bidirectional RNA transport along microtubules. Fusing CIBN to the minus-end-directed motor KIFC1* retrogradely transported *GAPDH* mRNA to the centrosome (22.3% at centrosome vs 1.9–5.4% controls; **Figure 1J**; **S2G**, **H**, **N**), while using the plus-end-directed motor KIF5B* drove the transcript anterogradely to the cell periphery (66.1% of cells showing enrichment; **Fig. 1K**; **S2G**, **I**, **O**) (*32*).

### Light-programmed RNA transport delivers second-scale, reversible kinetics

Opto-CRISPR-TO converts light into rapid and reversible control of RNA positioning in living cells. When targeting endogenous *GAPDH* mRNA to the OMM in HeLa cells, blue light induced enrichment that saturated within ∼30 minutes, substantially faster than chemical-based approaches (∼4 hours) (**Figure 2A**) (*19*). To resolve single-particle dynamics with higher temporal resolution, we used a U2OS reporter cell line expressing MS2-tagged transcripts and visualized their active transport via the motor protein KIFC1* (*19*, *38*). Upon blue-light stimulation (200 ms blue light on for every 5 s), transcripts began accumulating at the centrosome within 20 seconds, reaching saturation by ∼6 minutes (**Figure 2B**). Individual particles moved at 0.15 ± 0.02 µm·s⁻¹, a velocity consistent with known kinesin-driven transport (*39*). Trajectory analysis distinguished between sub-diffusive/diffusive and super-diffusive populations, with only the latter showing continuous, directed movement toward the centrosome.

**Figure 2.**
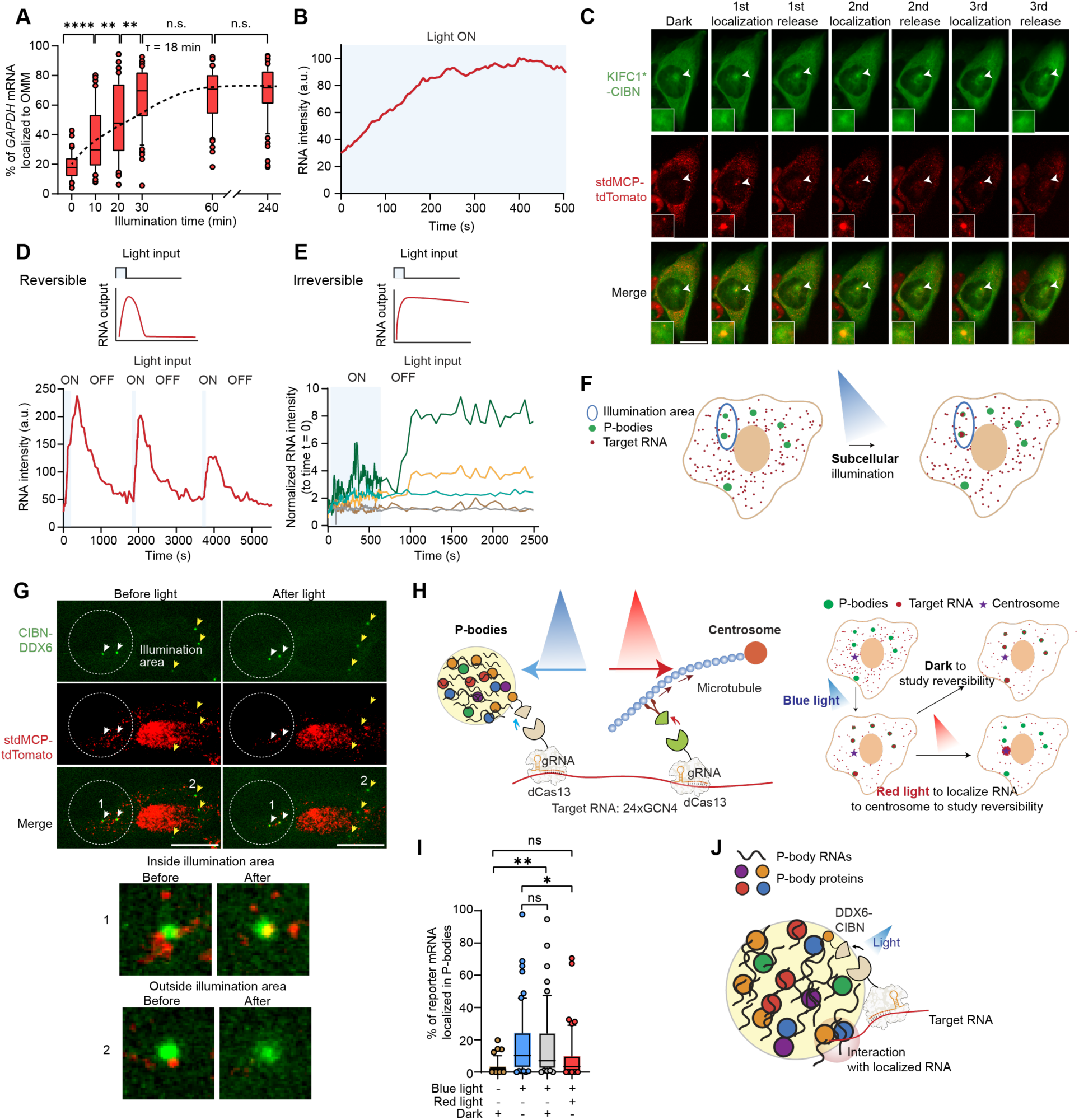
Opto-CRISPR-TO enables second-scale, reversible, multiplex RNA localization in cells with subcellular spatial resolution. (**A**) Quantification of *GAPDH* mRNA localization kinetics to the OMM during pulsed blue-light stimulation. From left to right, n = 62, 54, 41, 55, 53, 59 cells for each group. The dashed curve shows an exponential fit. (**B**) Time course of MS2-tagged reporter RNA fluorescence intensity at the centrosome for KIFC1*-mediated recruitment of MS2-tagged reporter mRNA to the centrosome. (**C**) Representative images showing KIFC1*-mediated recruitment of MS2-tagged reporter mRNA to the centrosome and subsequent release during three consecutive light/dark cycles. Scale bar: 20 µm. The insets show enlarged views of the centrosome. (**D**) Time course of reporter RNA fluorescence intensity at the centrosome for the cell shown in (C). (**E**) Representative traces of reporter RNA fluorescence intensity in five P-bodies before, during, and after blue-light stimulation, normalized to the initial intensity. Blue shading in (B), (D), and (E) indicates periods of pulsed blue-light stimulation. The top two P-bodies merged at ∼1000 s. (**F**) Schematic of spatially restricted RNA recruitment to P-bodies using local illumination within a single cell. (**G**) Representative images showing reporter RNA recruitment to P-bodies within the illuminated region (white circle), with enlarged views of P-bodies inside and outside this region. Scale bar: 20 µm. (**H**) Schematic of multiplex RNA control: blue light recruits 24×*GCN4* reporter RNA to P-bodies, followed by red-light-induced motor transport toward the centrosome. (**I**) Percentage of 24×*GCN4* reporter RNA in P-bodies before illumination, after 1 h of blue light, and after an additional 30 min of darkness or red light. From left to right, n = 49, 60, 54, 52 cells for each group. (**J**) Model illustrating potential interactions between recruited RNA and endogenous P-body components. *P* values were calculated using one-way ANOVA tests followed by Tukey’s HSD post-hoc tests for (A) and (I). n.s., *P* ≥ 0.05; \**P* < 0.05; \*\**P* < 0.01; \*\*\*\**P* < 0.0001. Boxplots show the median, 25th–75th percentiles (boxes), 10th–90th percentiles (whiskers), and individual values beyond the whiskers. 0 min, 60 min data in (**A**) was also used in Fig. 1C.

The system is also highly reversible. During repeated light/dark cycles, RNA accumulated and then dispersed with a ∼4-minute delay post-illumination, corresponding to the known dissociation half-life of the CRY2-CIBN interaction (t½∼5.5 min) (**Figure 2C-D**) (*24*). We demonstrated three consecutive cycles of recruitment and release, showcasing a temporal resolution not achievable with drug-based systems.

### Subcellular addressability reveals condensate-dependent control of RNA fate

We next tested if optogenetic routing could control RNA flux into and out of biomolecular condensates in live cells. Using DDX6-tethered Opto-CRISPR-TO, blue light rapidly recruited MS2-tagged transcripts into P-bodies (**Figure 2E**). Crucially, unlike the rapid dispersal seen from centrosomes (**Figure 2C-D**), RNAs routed to P-bodies remained trapped for over 30 minutes after the light was turned off. This suggests the condensate’s biochemical environment stabilizes the RNA through interactions with endogenous components, effectively "locking" it in place. The P-bodies exhibited liquid-liquid phase separation properties (*40*). The amount of recruited transcripts correlated strongly with P-body size. The mobility of the condensates was largely independent of their size, suggesting tethering or other constraints on P-bodies (*41*).

Because addressability is intrinsic to light, patterned illumination enables RNA recruitment to a micro-scale region of interest (ROI). We induced transcript capture exclusively by P-bodies within that ROI, while adjacent condensates in the same cell remained unaffected (**Figure 2F-G**), demonstrating the ability of Opto-CRISPR-TO to precisely control RNA localization at the subcellular level.

To achieve multiplex control, we combined blue-light-induced sequestration with red-light-induced active transport. After trapping 24xGCN4 reporter RNA in P-bodies using blue light, subsequent activation of a red-light-sensitive motor protein (PhyB–KIFC1*) actively extracted the transcripts, significantly accelerating their release compared to passive dissociation in dark (**Figure 2H-I**). Intriguingly, we observed that some of the P-body anchor protein, DDX6, was co-transported with the RNA to the centrosome, suggesting that the motor protein extracts the entire ribonucleoprotein (RNP) complex, in which RNA interacts with other P-body components, after being forced to P-bodies (**Figure 2J**) (*42*). Together, these results establish Opto-CRISPR-TO as a programmable "flux valve" to control RNA entry into, retention within, and active extraction from condensates.

### Opto-CRISPR-TO enables programmable control of endogenous RNA localization in the retina

To test Opto-CRISPR-TO *in vivo*, we chose the mouse retina, a highly structured and polarized tissue where the functional consequences of RNA localization are largely unexplored but can be interrogated via electroretinography (*43–45*). Baseline analysis using in situ hybridization technology via RNAscope (*46*) revealed that endogenous *Actb* mRNA is unevenly distributed across the retinal layers: it is scarce in the soma of the outer nuclear layer (ONL) and the inner plexiform layer (IPL), but enriched in the photoreceptor segments, abundant in the inner nuclear layer (INL) and ganglion cell layer (GCL).

To deliver Opto-CRISPR-TO components into mouse retinal cells in vivo, we used the established *in vivo* electroporation technique (*47*). The Opto-CRISPR-TO components (dCas13-CRY2, CIBN-KIFC1*, and *Actb*-targeting gRNAs or gNT control) were electroporated into mouse retinas *in vivo* at postnatal day zero (P0) to achieve motor-driven retrograde transport of *Actb* mRNA (**Figure 3A-B**). At P21, when the retinas were developed, we conducted 4 hours of blue-light cycling and found that retinas from the gT group displayed robust *Actb* mRNA relocalization, forming large perinuclear clusters, while the mRNA distribution remained undisturbed in both light-treated gNT and dark-treated gT controls (**Figure 3C**). This light- and guide-dependent phenotype was highly penetrant (observed in 5 of 6 treated mice vs. 0 of 14 controls) and disrupted the normal distribution of *Actb* mRNA across the retinal layers (**Figure 3C-D**).

**Figure 3.**
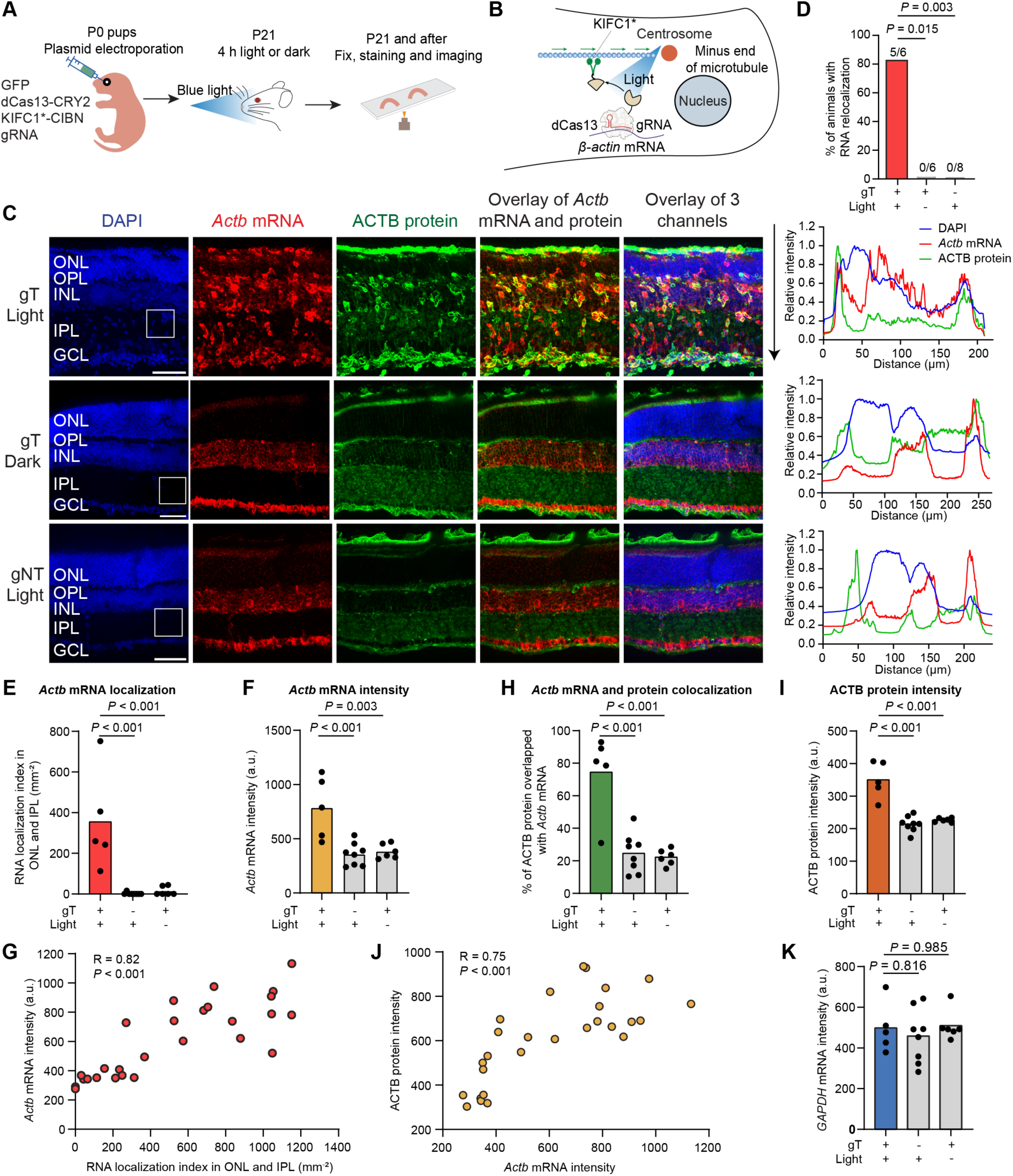
Opto-CRISPR-TO enables RNA mislocalization in the mouse retina *in vivo*. (**A**) Experimental timeline showing retinal electroporation at P0, followed by 4 h of pulsed blue light or darkness at P21. Retinas were harvested immediately after light treatment for fixation and processing. (**B**) Schematic showing light-induced, KIFC1*-mediated transport of *Actb* mRNA toward microtubule minus ends. (**C**) Representative retinal sections showing DAPI, *Actb* mRNA, and ACTB protein with corresponding relative intensity profiles across retinal layers after 4 h light treatment. Image contrast was adjusted individually to visualize spatial distributions. Scale bar: 50 µm. (**D**) Percentage of animals with RNA relocalization; animal counts are indicated above the bars. (**E**) *Actb* mRNA localization index (cluster density, mm⁻²) in the ONL and IPL. (**F**) Statistical analysis of *Actb* mRNA fluorescence intensity across retinal sections. (**G**) Correlation between the RNA localization index and *Actb* mRNA intensity. (**H**) Percentage of ACTB protein colocalized with *Actb* mRNA in the retina. (**I**) Statistical analysis of ACTB protein fluorescence intensity across retinal sections. (**J**) Correlation between *Actb* mRNA and ACTB protein intensities. (**K**) Statistical analysis of *Gapdh* mRNA intensity across retinal sections. For (G) and (J), n = 27 retinal regions from 4 animals; R, Pearson correlation coefficient. *P*, corresponding *P* value. For (E), (F), (H), (I), and (K), each point represents one animal and bars indicate means; n = 5, 8, and 6 animals for gT + light, gNT + light, and gT + dark, respectively. *P* values were calculated using two-sided Fisher’s exact tests (D) or one-way ANOVA followed by Tukey’s HSD post hoc tests (E, F, H, I, and K). ONL/INL, outer/inner nuclear layer; OPL/IPL, outer/inner plexiform layer; GCL, ganglion cell layer.

To quantify mRNA relocalization in mouse retina *in vivo*, we calculated a ‘localization index’ (density of *Actb* mRNA clusters) in the ONL and IPL, where baseline *Actb* mRNA levels are low. The localization index was dramatically increased in retinas treated with Opto-CRISPR-TO (354.4 mm⁻^2^) compared to gNT (2.0 mm⁻^2^) and dark (14.1 mm⁻^2^) controls (**Figure 3E**). These results confirm the robustness of Opto-CRISPR-TO in repositioning endogenous mRNA *in vivo*.

### *In vivo* RNA relocalization promotes transcript accumulation

To test whether spatial repositioning alters transcript levels, we quantified *Actb* mRNA intensity after Opto-CRISPR-TO perturbation. Strikingly, retinas with guide-targeted, light-induced *Actb* mRNA relocalization showed significantly higher whole-retina *Actb* mRNA signal than controls (gNT + light or gT + dark; **Figure 3F**). Across retinal regions, this increase in intensity strongly correlated with the degree of local RNA clustering, as calculated by the localization index (R = 0.82), indicating a plausible causal link between enforced perinuclear *Actb* mRNA relocalization and higher steady-state transcript levels *in vivo* (**Figure 3G**). The effect was target-specific, as the pattern and intensity of *Gapdh* mRNA were unchanged across conditions (**Figure 3K**; **S3**).

Cell-type co-staining with *Actb* mRNA revealed that the *Actb* mRNA relocalization and subsequent increase in intensity occurred across diverse retinal cells, including photoreceptors (∼32% of total relocalization signals) (*48*), bipolar cells (∼21%) (*49*), Müller glia (∼7%) (*50*), and amacrine cells (∼5%) (*51*). Thus, light-programmed RNA positioning broadly alters the target RNA intensity, likely by deviating it from degradation machineries (*52*, *53*), across multiple cell types in a multi-cellular mammalian tissue *in vivo*.

### RNA relocalization reprograms local protein translation *in vivo*

We next investigated whether relocalizing *Actb* mRNA altered its translation *in vivo*. In untreated retina, *Actb* mRNA and ACTB protein only partially overlap. For example, both *Actb* mRNA and ACTB protein enrich in photoreceptor segments, but INL shows high *Actb* mRNA with low protein, and IPL shows low *Actb* mRNA with strong ACTB protein, consistent with spatially decoupled translation *in vivo* (*54*).

After 4-hour Opto-CRISPR-TO treatment, cells with forced *Actb* mRNA relocalization also showed robust ACTB protein enrichment within the same perinuclear clusters, while controls retained wildtype-like patterns (**Figure 3C**). This demonstrates that the repositioned mRNA is actively translated at its new site. As a result, the relocated RNA and protein were now highly co-localized (**Figure 3H**). This also indicates that precise RNA localization is crucial for ensuring correct protein distribution, as ACTB proteins translated from *Actb* mRNAs localized at the soma were enriched there and not efficiently delivered to their native location. In addition, the total intensity of ACTB protein significantly increased (**Figure 3I**), which is strongly correlated with the increase in RNA intensity (R = 0.75; **Figure 3J**). Thus, forced RNA positioning acutely elevates and redirects local translation and protein localization in the retina *in vivo*.

### Short-term RNA relocalization causes lasting retinal dysfunction

Given the significant local changes in mRNA and protein levels, we next investigated whether brief, 4-hour relocalization of *Actb* mRNA could lead to retinal dysfunction. We used electroretinography (ERG) to measure the electrical responses of the retina to light, where the a-wave and b-wave report the function of photoreceptors (scotopic: rods; photopic: cones) and bipolar cells, respectively (*55*, *56*). The mice, which were electroporated at P0 (gT or gNT) and received 4-hour blue light at P21, were dark-adapted overnight and then tested for ERG; only eyes with high electroporation efficiency were analyzed (**Figure 4A**). ERGs were recorded under scotopic (dark-adapted) conditions (0.01, 0.1, 1 cd·s·m^-^²) and photopic (light-adapted) conditions (after 10-min light adaptation; 3, 10 cd·s·m^-^²).

**Figure 4.**
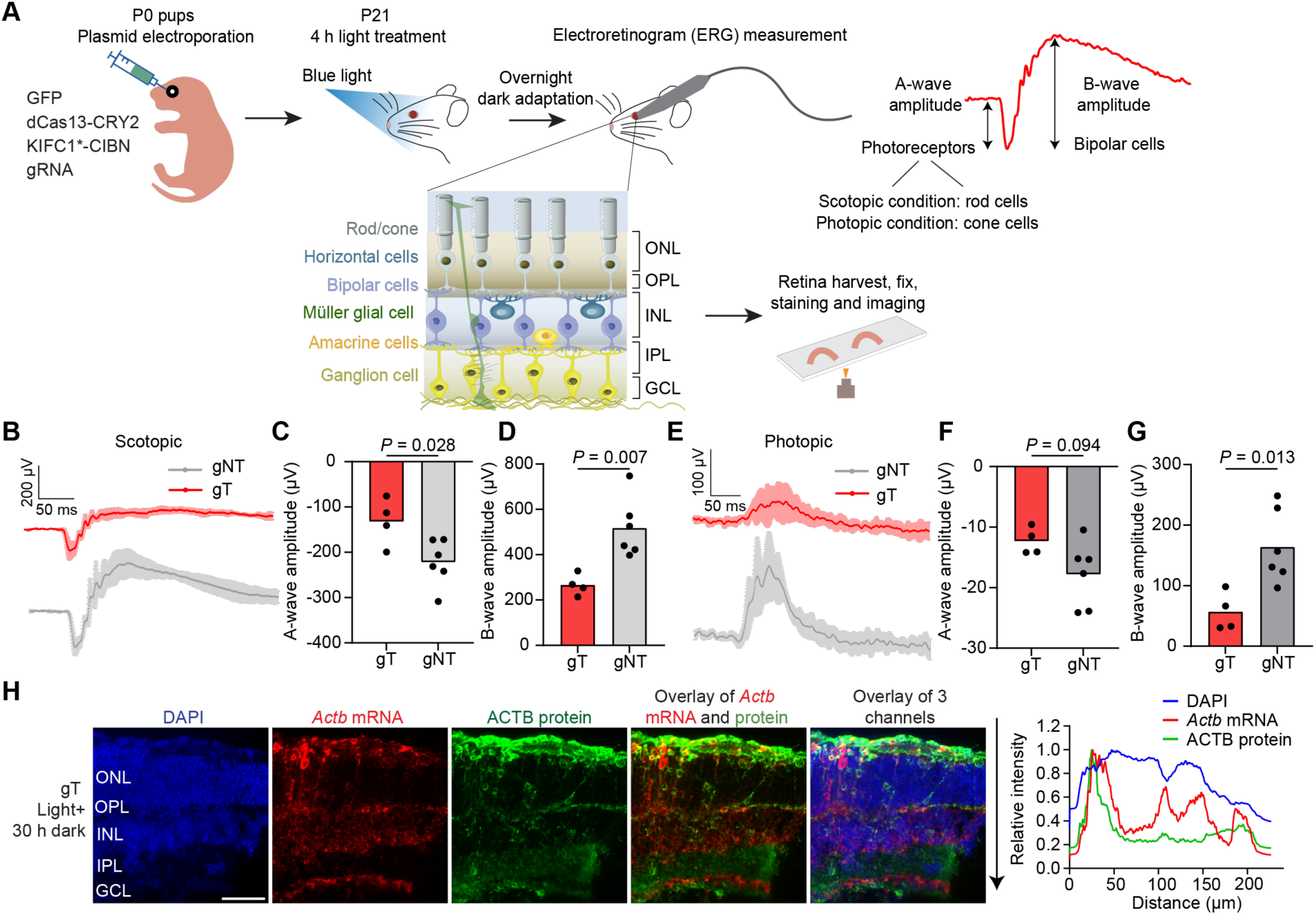
Acutely induced RNA mislocalization causes persistent RNA perturbation and impairs visual function. (**A**) Schematic of experimental procedure for electroretinogram (ERG) measurements: retinal electroporation of Opto-CRISPR-TO components at P0 pups, 4 h of pulsed blue light or darkness at P21, overnight dark adaptation, ERG recording, and tissue collection for fixation and processing. (**B–D**) Scotopic ERG traces at 1 cd·s·m⁻² (**B**) and corresponding statistical analysis of a-wave (**C**) and b-wave (**D**) amplitudes. (**E–G**) Photopic ERG traces at 10 cd·s·m⁻² after 10 min of light adaptation (**E**) and corresponding statistical analysis of a-wave (**F**) and b-wave (**G**) amplitudes. Traces show mean ± SD. (**H**) Representative retinal section for n = 4 animals showing DAPI, *Actb* mRNA, and ACTB protein distribution after 4 h of pulsed blue light followed by 30 h of darkness, with corresponding relative intensity profiles across retinal layers. Scale bar: 50 µm. For (B–G), n = 4 and 6 for gT + light and gNT + light, respectively. In (C), (D), (F), and (G), each point represents one animal and bars indicate means. *P* values were calculated using two-tailed unpaired t-tests.

Under scotopic conditions, both a-wave and b-wave amplitudes were significantly reduced in the targeted group compared to controls, indicating impaired function of both rod and bipolar cells following *Actb* mRNA relocalization (**Figure 4B-D**; **S4**). Under photopic conditions, a-waves were unaffected, consistent with the observation that *in vivo* electroporation at P0 is unable to deliver plasmid DNA into cone cells, whereas b-waves were significantly reduced in the targeted group, confirming a bipolar cell deficit across lighting regimes (**Figure 4E-G**; **S4**).

These results demonstrate that a transient, optogenetically driven relocalization of a single endogenous mRNA is sufficient to produce a specific and lasting functional impairment of the retinal circuit.

### Transient RNA relocalization induces lasting changes in cell positioning

We next investigated the cellular mechanisms underlying retinal dysfunction following transient RNA relocalization. The most dramatic consequence of 4-hour *Actb* mRNA relocalization was a profound disruption of retinal architecture. We observed a significant increase in cell nuclei within the IPL, a region normally devoid of cell bodies (**Figure 5A-B**). Most of these misplaced cells (∼71%) had *Actb* mRNA repositioning, directly linking the molecular event to the cellular phenotype (**Figure 5C**). This change in cell position is consistent with the essential role of β-actin in maintaining cytoskeletal structure and cell motility (*57*).

**Figure 5.**
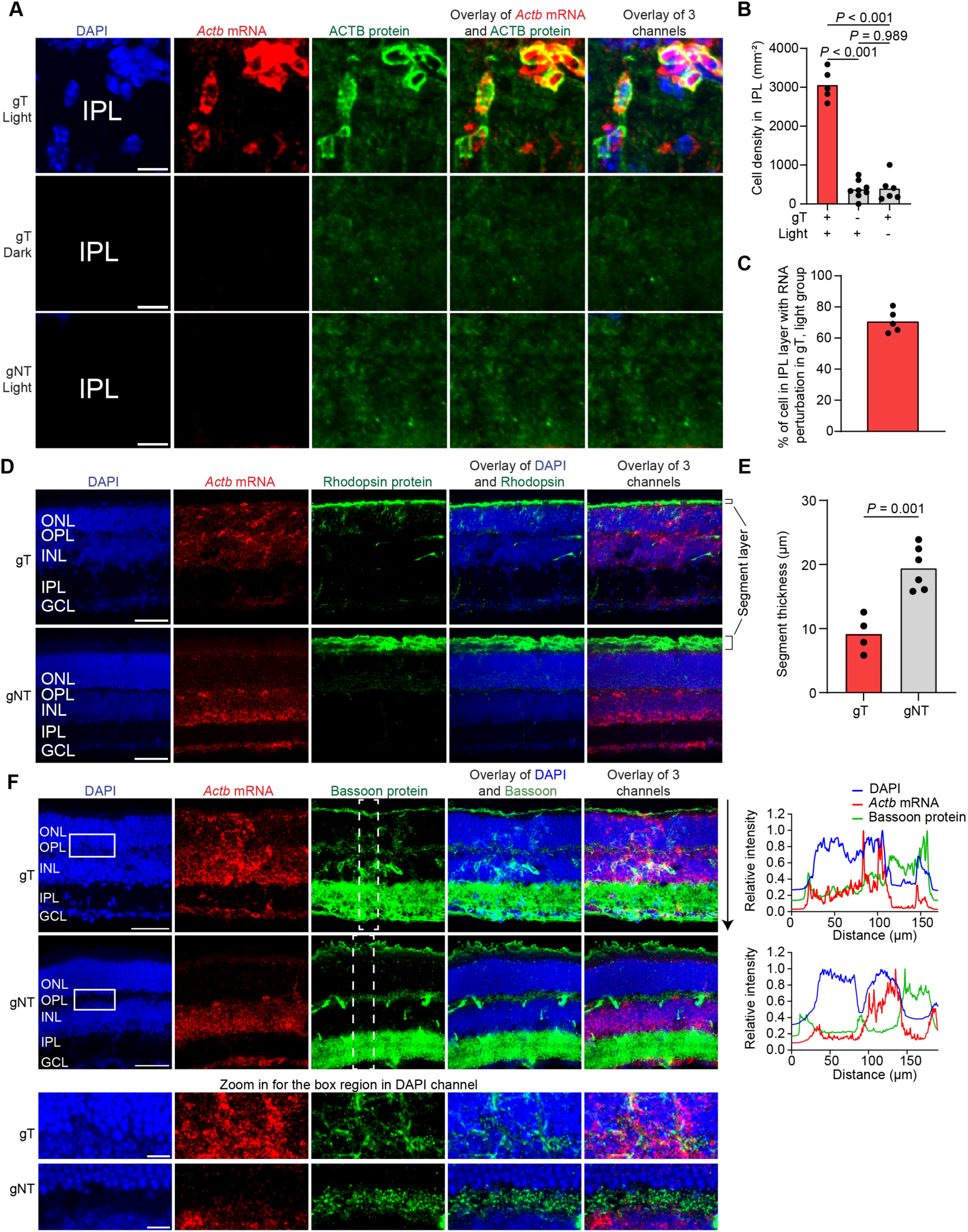
RNA mislocalization disrupts retinal structure. (**A**) Representative retinal slices showing DAPI, *Actb* mRNA, and ACTB protein in the IPL region, with the insert magnified in Figure 3C. Scale bar: 10 µm. (**B**) Cell nuclei density in IPL region across groups. From left to right, the number of animals is 5, 8, 6. Each data point and the mean are shown. *P* values were calculated using one-way ANOVA tests followed by Tukey’s HSD post-hoc tests. (**C**) Percentage of cell nucleus in IPL region with RNA perturbation for the gT, light group. The number of animals is 5. Each data point and the mean are shown. (**D**) Representative retinal slices showing DAPI, *Actb* mRNA, and Rhodopsin protein for the gT and gNT groups after 4 h of light and overnight dark treatment, and ERG measurement. (**E**) Thickness of the segment layer for gT and gNT groups after 4 h light, overnight dark treatment, and ERG measurement. From left to right, the number of animals is 4 and 6. Each data point and the mean are shown. *P* value was calculated using a two-tailed unpaired t-test. (**F**) Representative retinal slices showing DAPI, *Actb* mRNA, and Bassoon protein in the gT and gNT groups after 4 h of light exposure, overnight dark treatment, and ERG measurement. The relative average intensity for the rectangular region on the Bassoon channel for gT and gNT groups was plotted. Scale bar: 50 µm. The lower two rows magnify the white rectangular region on the DAPI channel.

We next asked if this cellular reorganization was reversible. In stark contrast to the rapid RNA dispersal observed in vitro upon light withdrawal (**Figure 2C-D**), the *in vivo* changes were prolonged throughout retina development. Mice examined 30 hours after the initial 4-hour light pulse still exhibited dramatic cell mis-positioning and disrupted ACTB protein distribution (**Figure 4H**). This suggests that short-term, molecular-level RNA relocalization can trigger an extended pathological cascade.

### Synaptic and photoreceptor architecture is disrupted without apoptosis

To further probe mechanisms underlying the ERG deficits, we examined photoreceptor structure and synaptic organization. Rhodopsin-positive segments were significantly shortened after *Actb* mRNA relocalization (**Figure 5D-E**) (*58*, *59*), consistent with reduced scotopic a-waves and indicating compromised rod phototransduction. Bassoon, a scaffold of ribbon synapses that transmit signal from photoreceptor cells to bipolar cells (*60*), was disrupted in targeted, light-treated retinas compared to gNT controls (**Figure 5F**), providing an anatomical correlation of impaired bipolar-cell b-waves.

Beyond the synapse, we observed a profound loss of tissue architecture. Phalloidin staining revealed marked F-actin disorganization (*61*), with loss of the characteristic enrichment in the segment and OPL, accompanied by disturbed overall histology (**Figure S5**). These changes support a cytoskeletal mechanism that links RNA mislocalization to altered cell positioning and synaptic structure.

Importantly, this widespread dysfunction was not due to cell death, as the level of the apoptosis marker Cleaved Caspase-3 was not significantly increased in treated retinas compared with the gNT group (**Figure S6**) (*62*).

## DISCUSSION

This work establishes Opto-CRISPR-TO as a universal, *in vivo*-ready method to manipulate endogenous RNA geography with high temporal resolution (second-scale control) and subcellular spatial resolution. By using light to recruit dCas13-bound transcripts to defined compartments or along directed transport routes, we convert RNA mislocalization from a correlation into a causal variable. A brief mislocalization of a single mRNA (*Actb*) induces prolonged changes in protein distribution, synaptic architecture, cell positioning, and organ function, providing experimental evidence that RNA mislocalization causes pathological effects.

Methodologically, optogenetic induction confers speed, reversibility, and subcellular addressability without systemic drugs, enabling within-tissue perturbations and multiplex control in the same cell. Coupling optogenetic control to CRISPR-TO enables targeting theoretically any endogenous RNA (mRNA or noncoding RNA) in living cells.

Biologically, the approach demonstrates that RNA position governs local translation and downstream cell biology. In the retina, transient *Actb* mRNA relocalization (hours) increased local RNA and protein intensities, reorganized F-actin, shortened rhodopsin-rich outer segments, and disrupted Bassoon-positive ribbon synapses and cell positioning without inducing apoptosis (**Figure 6**). These changes explain the mouse vision deficits we observed in ERG tests. Thus, local transcript distribution and availability are sufficient to dramatically remodel morphology, connectivity, and physiology.

**Figure 6.**
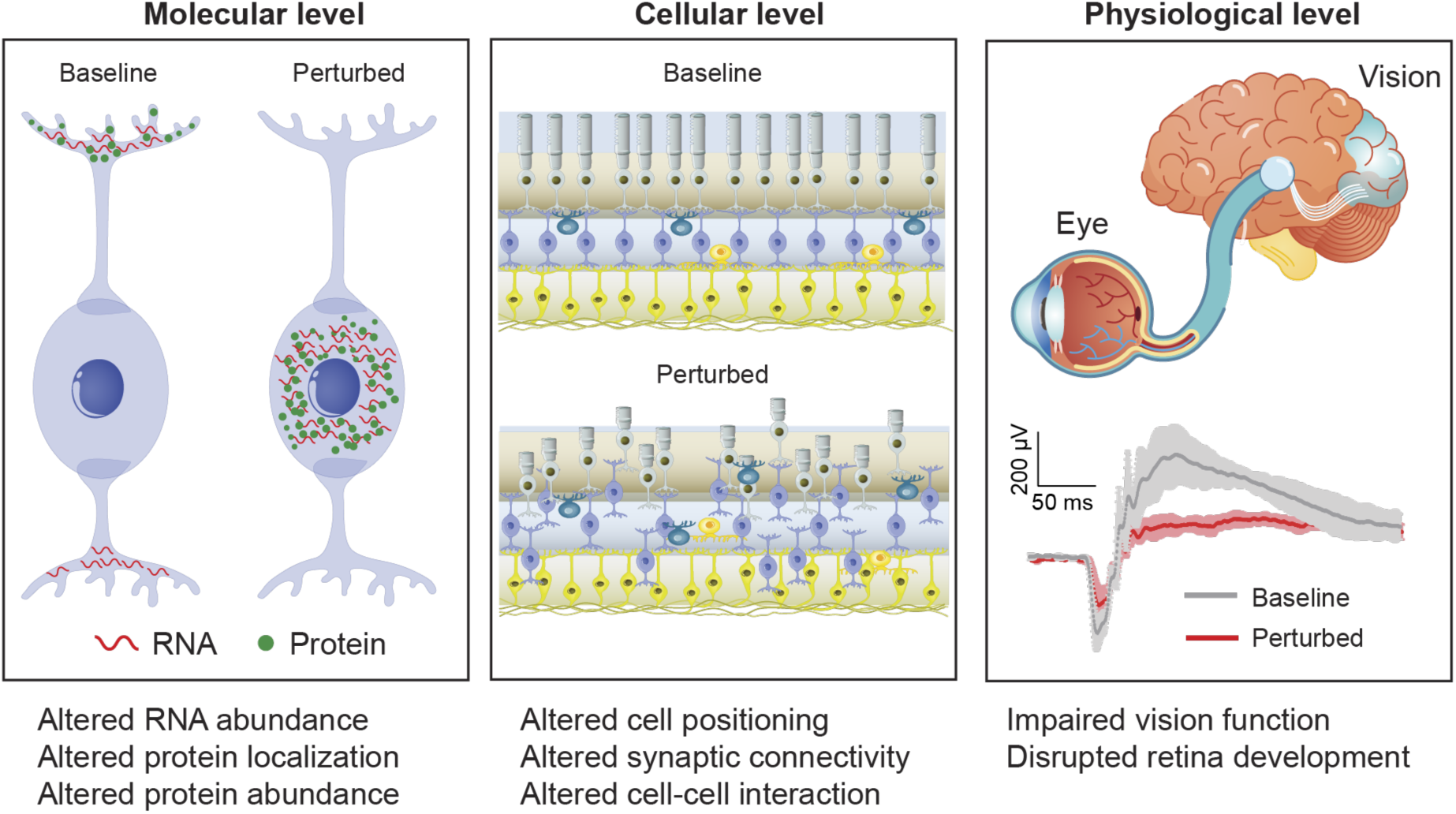
Schematic summary of the molecular, cellular, and physiological consequences of RNA mislocalization.

Our observations support a “local abundance” paradigm: cellular phenotypes depend not just on the total amount of a specific RNA but also on its distribution relative to compartments, organelles, motors, and synapses. Pathology can arise from local RNA imbalance (too much or too little local RNA at an important site such as synapses). This framework aligns with clinical genetics in neurodevelopment and neurodegeneration, where RNA-granule dynamics and axonal transport are aberrant (*12, 15*); the potential causal relationship between RNA mislocalization and these pathologies is now testable. Further uses of Opto-CRISPR-TO may (i) rescue disease models by correcting localization; (ii) map dose-location-response landscape to identify therapeutic “sweet spots” across various RNAs; and (iii) multiplex RNA localizations towards multiple destinations to decode “combinatorial RNA geography” underlying resilience or vulnerability.

We focused on *Actb* mRNA and the retina, which were chosen for their polarization, optical access, and tractable delivery. Future work could integrate ribosome tagging or nascent-peptide reporters to read out compartment-specific translation in real time, combine with single-cell transcriptomics and electrophysiology to link spatial control to circuit output, and expand delivery formats such as adeno-associated virus (AAV) vectors using compact proteins for broader tissue applications. Finally, correcting mislocalization rather than transcription may suffice to restore function in disorders in which RNA is present but misplaced.

In summary, controlling where RNAs live in cells reveals and enables intervention in “the spatial logic of RNA regulation” in living organisms. Opto-CRISPR-TO provides a causal approach to connect RNA organization to protein synthesis, cell architecture, and organ physiology, opening a new path toward therapies that normalize RNA location rather than RNA levels.

## Acknowledgments

We thank all members of the Lei Stanley Qi lab for facilitating the experiments and for useful discussions. We thank Prof. W.E. Moerner’s lab for providing Phalloidin Labeling Probes. We thank Dr. David L. Spector at Cold Spring Harbor Laboratory for U2OS 2-6-3 cells. We thank Stanford University Cell Sciences Imaging Core Facility (RRID:SCR_017787) for assisting with microscopy imaging.

## Author contributions

S.C. and L.S.Q. conceived the study. S.C., W.C., G.H., S.W., and L.S.Q. designed the experiments. S.C., B.D., M.L.F., A.A., P.P., G.W., A.C., D.H., M.H., and Y.S. performed the experiments. S.C., A.A., A.C., D.H., M.H., Y.Z., X.W., and L.S.Q. analyzed the data. S.C. and L.S.Q. prepared the figures and wrote the original manuscript draft. All authors reviewed and edited the manuscript. L.S.Q. secured funding.

## METHODS

### Cell culture

HEK293T (ATCC, CRL-3216), HeLa (Sigma, 93021013), and U2OS 2-6-3 cell lines (a gift from Dr. David L. Spector, Cold Spring Harbor Laboratory) were cultured in DMEM medium (Thermo Fisher, 10569-044) supplemented with 10% FBS (Sigma-Aldrich, F0926). Neuro-2a cells (CCL-131, ATCC) were cultured in EMEM (30-2003, ATCC) with 10% FBS. Cortical neurons from C57BL/6J (The Jackson Laboratory, 000664) P0 pups were dissected and maintained as described in the reported protocol (*19*). All cells were cultured at 37 °C in incubators with 5% CO_2_.

### Animals

Wild-type neonatal mice obtained from timed pregnant CD-1 mice (Charles River Laboratories, #022) were used for *in vivo* electroporation experiments. Animals of mixed sex, as obtained from the supplier without further stratification, were used in equal or random proportions. All animal studies were approved by the Institutional Animal Care and Use Committee at Stanford School of Medicine. C57BL/6J (000664, The Jackson Laboratory) P0 pups were used for collecting cortical neurons. All animal procedures were approved by Stanford University’s Administrative Panel on Laboratory Animal Care. Animals were maintained on a 12 h light-12 h dark cycle at 20–25 °C temperature and 50–65% humidity in Stanford University’s Veterinary Service Center (VSC) and fed with food and water ad libitum as appropriate.

### Imaging microscopes

Confocal images were taken at Stanford University Cell Sciences Imaging Facility using a Nikon Ti2 Crest SDC microscope. Widefield images were taken using a Leica dMi8 microscope. For retina samples, images were taken with the Nikon Ti2 Crest SDC microscope, with Z-stacks acquired at a 1 μm step size using a 40x S Fluor oil objective (NA = 1.3). For primary neuron samples, images were acquired using the Nikon Ti2 Crest SDC microscope, producing Z-stacks with a 0.3 μm step size and a 40x S Fluor oil objective (NA = 1.3). For cell line samples, images were taken using the Nikon Ti2 Crest SDC microscope with Z-stacks at a 0.3 μm step size using a 60x Plan Apo oil objective (NA = 1.40) or the Leica dMi8 microscope with Z-stacks at a 0.3 μm step size using a HC PL APO 63x oil objective (NA = 1.40). H&E images were taken at Stanford University Cell Sciences Imaging Facility using a Nikon Ti widefield microscope with the 20x air objective (NA = 0.8).

### RNA-FISH and immunostaining

For HeLa, N2a, and HEK293T cells, cells were washed with 1x PBS, fixed with 4% PFA for 10 min at room temperature, and quenched with 50 mM glycine at room temperature. Cells were washed twice with 1x PBS and permeabilized in 70% ethanol for 1 h to overnight at 4 °C. Cells were then incubated in 2x SSC solution with 10% formamide for 5 min at room temperature.

Fixed and permeabilized cells were incubated in hybridization solution (0.12 μM FISH probes in Stellaris RNA FISH Hybridization Buffer (Stellaris, SMF-HB1-10) with 10% formamide) for 4 h to overnight at 37 °C. Cells were incubated in 2xSSC solution with 15% formamide for 30 min at 37 °C twice, washed with PBST (1x PBS with 0.1% Tween-20), washed with 2xSSC, and mounted on glass slides using VECTASHIELD PLUS Antifade Mounting Medium (Vector Laboratories, H-1900-10).

For cells with RNA FISH and antibody co-staining, cells were washed with 1x PBS, fixed with 4% PFA for 10 min at room temperature, and quenched with 50 mM glycine at room temperature. Cells were washed twice with 1x PBS and permeabilized with 0.1% TritonX-100 in 1x PBS containing 1 mM MgCl_2_ and 0.1 mM CaCl_2_ for 15 min at room temperature. Cells were incubated with primary antibody in PBS containing RNase Inhibitor (New England Biolabs, M0314S) for 60 min at 37 °C and washed with 1x PBS. Cells were then incubated in secondary antibody in PBS with RNase Inhibitor (New England Biolabs, M0314S) for 30 min at 37 °C and washed with PBST three times. Cells were then incubated in 2xSSC solution with 10% formamide for 10 min at room temperature. Cells were then incubated in hybridization solution and processed using the same protocol for cell FISH.

### Neuron FISH and immunostaining

Primary neurons were washed with 1x PBS, fixed with 4% PFA with 4% sucrose, 1 mM MgCl_2_, and 0.1 mM CaCl_2_ for 20 min at room temperature, quenched by 50 mM glycine at room temperature, and permeabilized with 0.1% TritonX-100 in 1xPBS with 1 mM MgCl_2_, and 0.1 mM CaCl_2_ for 15 min at room temperature. Neurons were incubated in 1:500 FITC anti-HA antibody (Genscript, A01621) in PBS with RNase Inhibitor (New England Biolabs, M0314S) for 30 min at 37 °C, then washed with 1x PBS. Neurons were then incubated in 2x SSC solution containing 10% formamide for 30 min at room temperature. Neurons were then incubated in hybridization solution and processed using the same protocol for cell FISH.

### Image analysis

MATLAB, ImageJ, Origin, Python, and GraphPad Prism software were used for image processing and analysis. Only transfected cells were counted in analysis.

### Electroretinogram (ERG) measurement

Mice were dark-adapted overnight and then anesthetized using 0.01 mg/g xylazine and 0.08 mg/g ketamine. Tropicamide ophthalmic solution (Sandoz, 61314-354-01) was applied to the mouse eye to induce dilation. Mice were then transferred to a Celeris visual electrophysiology system (Diagnosys, Celeris) for electroretinogram measurement. For scotopic measurement, single-flash ERGs were recorded after dark adaptation at luminances of 0.01, 0.1, and 1 cd·s/m². For photopic measurements, the animal was light-adapted for 10 min, and single-flash ERGs were recorded at luminances of 3 and 10 cd·s/m².

### Calculation of RNA localization index

RNA localization index is defined as the number of clusters (50 µm^2^ < cluster size < 1200 µm^2^) per unit area (mm^-2^). Clusters are identified using a centroid-based clustering method in MATLAB.

### Quantification of RNA intensity at centrosome

Custom MATLAB code was used to locate the centrosome center in each frame by selecting the brightest pixel. A circular ROI with a radius of 1 µm or 2 µm was drawn around the centrosome site, and the average RNA intensity within the ROI was measured.

### Statistics

For *in vitro* cell experiments, n indicates the number of cells or nuclei, and *P* values describe differences among cells from different groups. For *in vivo* quantification, unless otherwise noted, the unit of analysis was the animal, with one image analyzed per animal. Exact n values, *P* values, and statistical tests are provided in the figures and figure legends; exceptions are noted in the legends.

## SI Figures

**SI Figure 1.**
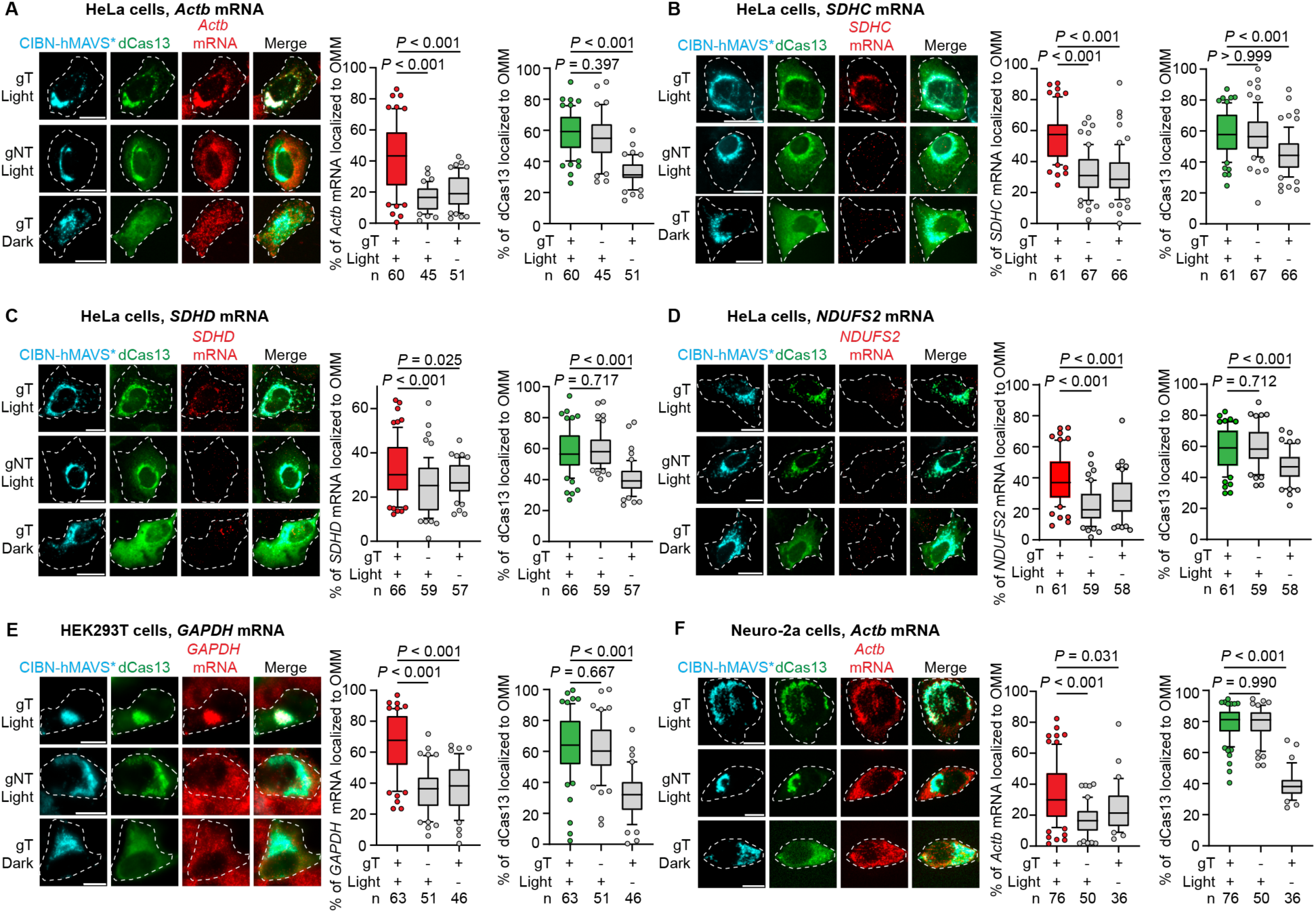
Opto-CRISPR-TO localizes distinct RNAs across cell types. (**A**-**D**) Left: representative images showing endogenous *Actb*, *SDHC*, *SDHD*, *NDUFS2* mRNA and dCas13 protein localization in HeLa cells. White dotted lines indicate cell shape. Scale bar: 20 µm. Right: percentage of targeted mRNA and dCas13 protein localized to the OMM for individual cells. (**E**) Representative images showing endogenous *GAPDH* mRNA and dCas13 protein localization in HEK293T cells. White dotted lines indicate cell shape. Scale bar: 10 µm. Right: percentage of *GAPDH* mRNA and dCas13 protein localized to the OMM for individual cells. (**F**) Representative images showing endogenous *Actb* mRNA and dCas13 protein localization in Neuro-2a cells. White dotted lines indicate cell shape. Scale bar: 10 µm. Right: percentage of *Actb* mRNA and dCas13 protein localized to the OMM for individual cells. *P* values were calculated using one-way ANOVA followed by Tukey’s HSD post-hoc tests for (A), (B), (C), (D), (E), (F). n is the number of cells in each group. Boxplots show the median, the 25–75% range (boxes), the 10–90% range (whiskers), and outliers.

**SI Figure 2.**
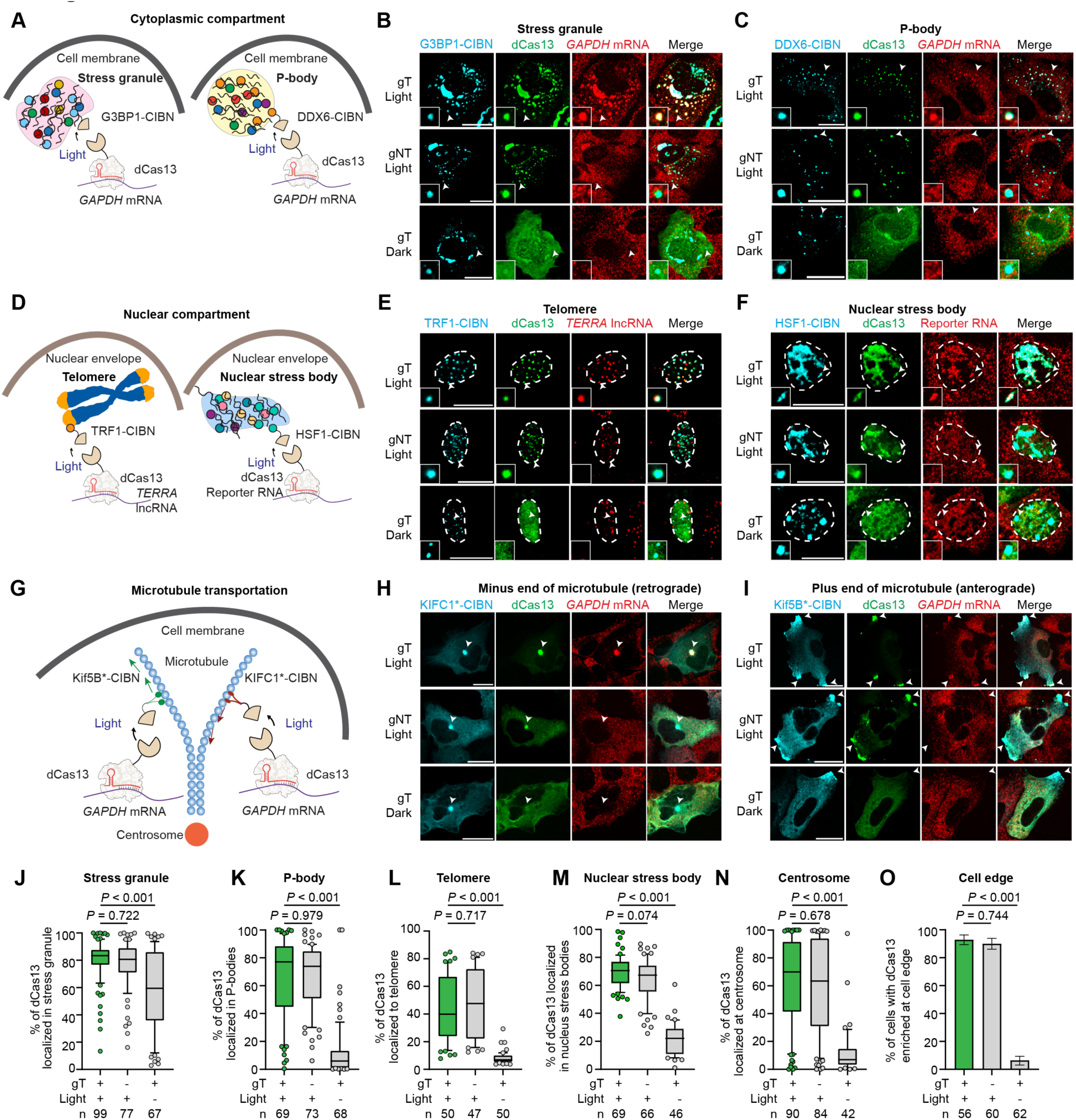
Opto-CRISPR-TO localizes different RNAs to various subcellular compartments. (**A**) Schematics of endogenous *GAPDH* mRNA targeting to stress granules and P-bodies. (**B, C**) Representative images showing endogenous *GAPDH* mRNA and dCas13 protein localization to (**B**) stress granules and (**C**) P-bodies. Insets magnify the regions indicated by arrows. (**D**) Schematics of endogenous *TERRA* long noncoding RNA targeting to telomeres and 24×*GCN4* reporter RNA targeting to nuclear stress bodies. (**E, F**) Representative images showing *TERRA* and dCas13 localization to telomeres (**E**), and 24×*GCN4* RNA and dCas13 localization to nuclear stress bodies (**F**). White dotted lines outline nuclei. Insets magnify the regions indicated by arrows. (**G**) Schematic of motor-driven *GAPDH* mRNA transport along microtubules. (**H, I**) Representative images showing *GAPDH* mRNA and dCas13 localization following minus-end-directed (**H**) or plus-end-directed (**I**) transport in HeLa cells. (**J–N**) Percentage of dCas13 localized to stress granules (**J**), P-bodies (**K**), telomeres (**L**), nuclear stress bodies (**M**), and centrosomes (**N**). Each data point represents one cell (**J, K, N**) or nucleus (**L, M**). (**O**) Percentage of transfected cells with dCas13 enriched at the cell periphery. *P* values were calculated using one-way ANOVA followed by Tukey’s HSD post-hoc tests (**J–N**) or pairwise two-sided Fisher’s exact tests without correction (**O**). Boxplots show the median, interquartile range, 10th–90th percentile whiskers, and points outside the whiskers. Scale bar: 20 µm. n indicates the number of cells or nuclei analyzed in each group.

**SI Figure 3.**
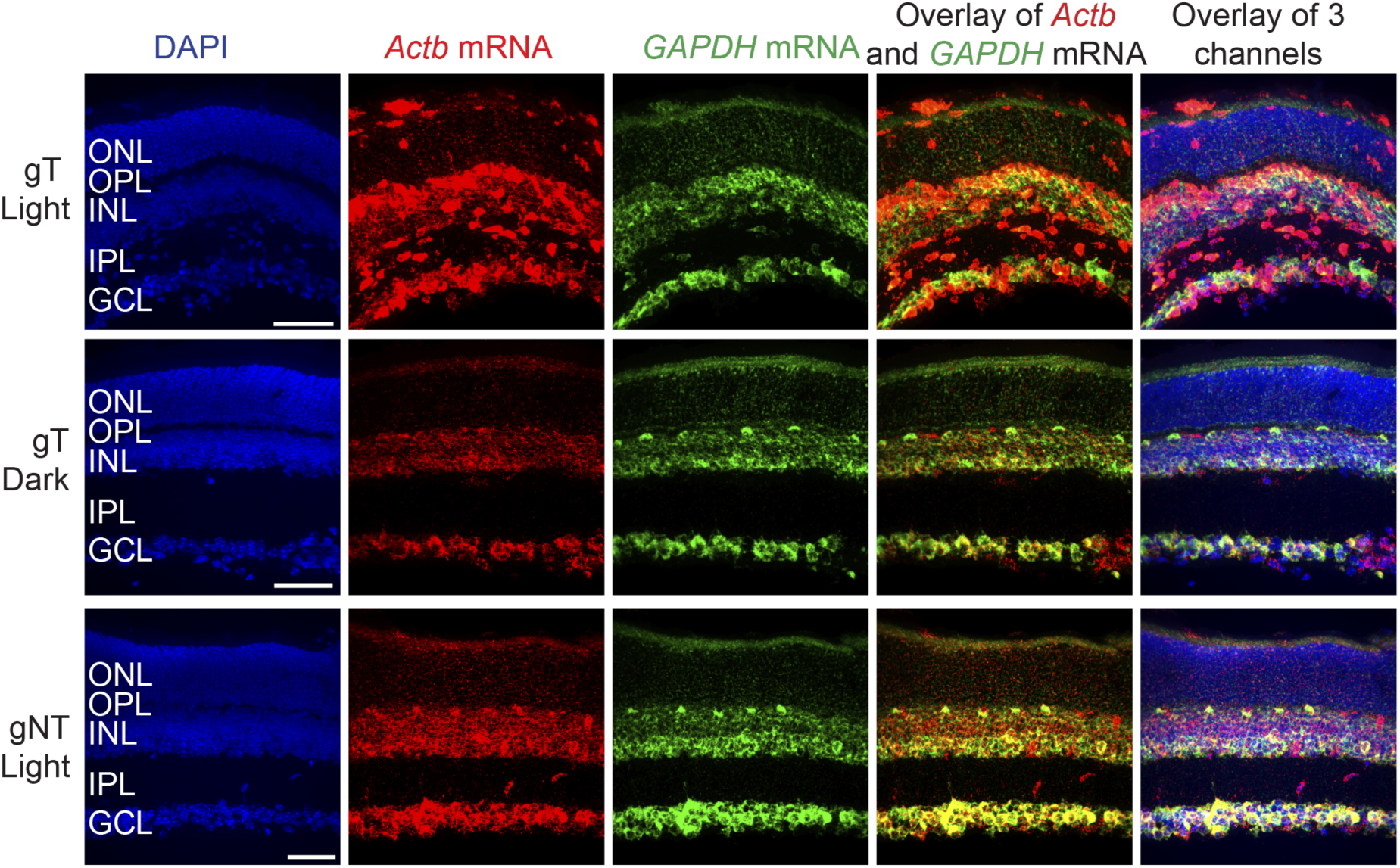
*Actb* and *Gapdh* mRNA distributions in retinal sections. Representative retinal sections showing DAPI, *Actb* mRNA, and *Gapdh* mRNA. For each RNA channel, identical contrast settings were applied across groups. Scale bar: 50 µm.

**SI Figure 4.**
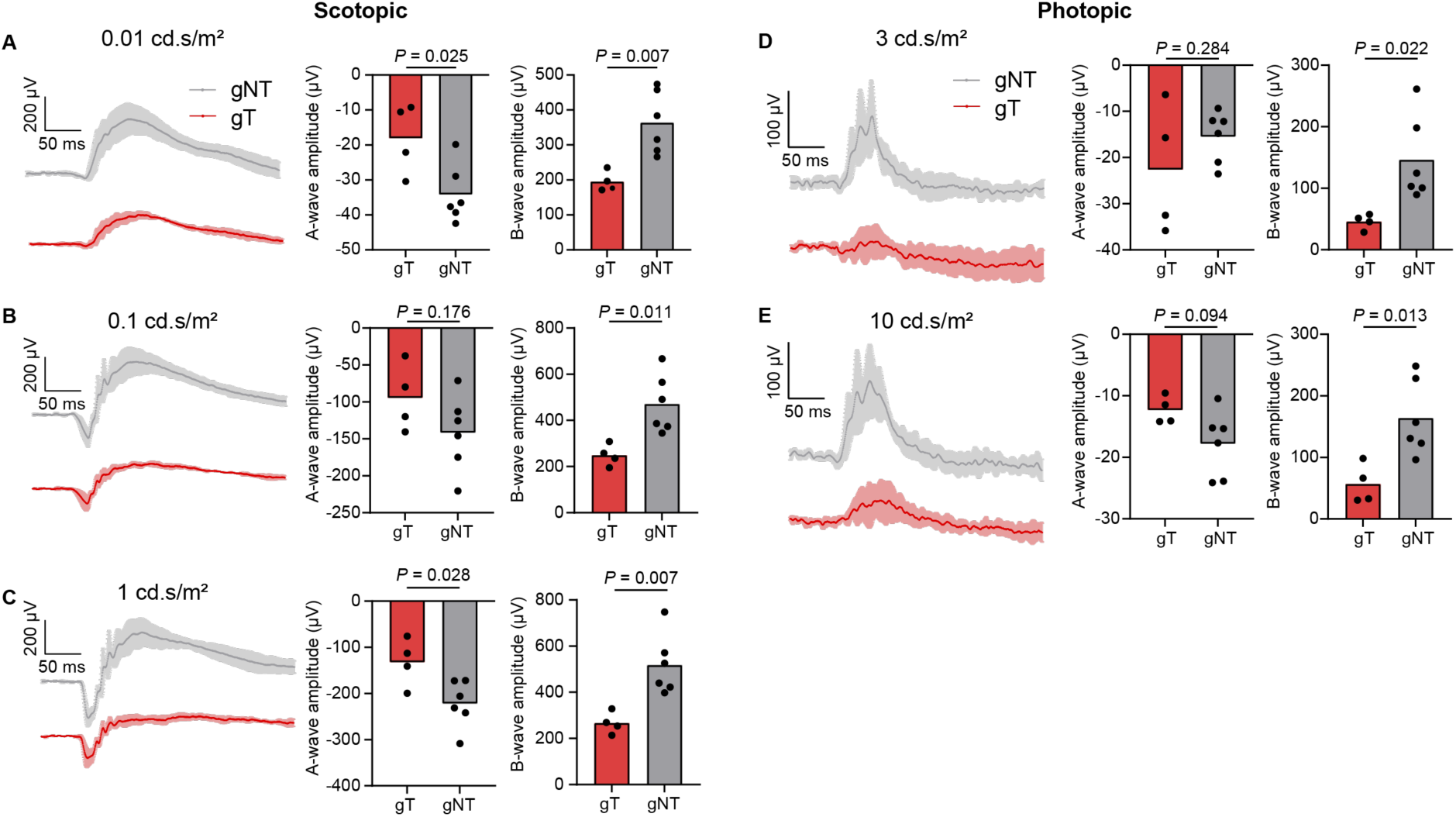
Electroretinographic responses after *Actb* mRNA relocalization. ERG measurement raw traces and statistical results of a-wave and b-wave under scotopic conditions with (**A**) 0.01 cds/m^2^ (**B**) 0. 1 cd.s/m^2^ (**C**) 1 cd.s/m^2^ light stimulus. ERG measurement raw traces and statistical results for the a-wave and b-wave under photopic conditions with (**D**) 3 cd.s/m^2^ (**E**) 10 cd.s/m^2^ light stimulus. From left to right, the numbers of animals are 4 and 6 across all plots. Each data point and the mean are shown. *P* values were calculated using a two-tailed unpaired t-test.

**SI Figure 5.**
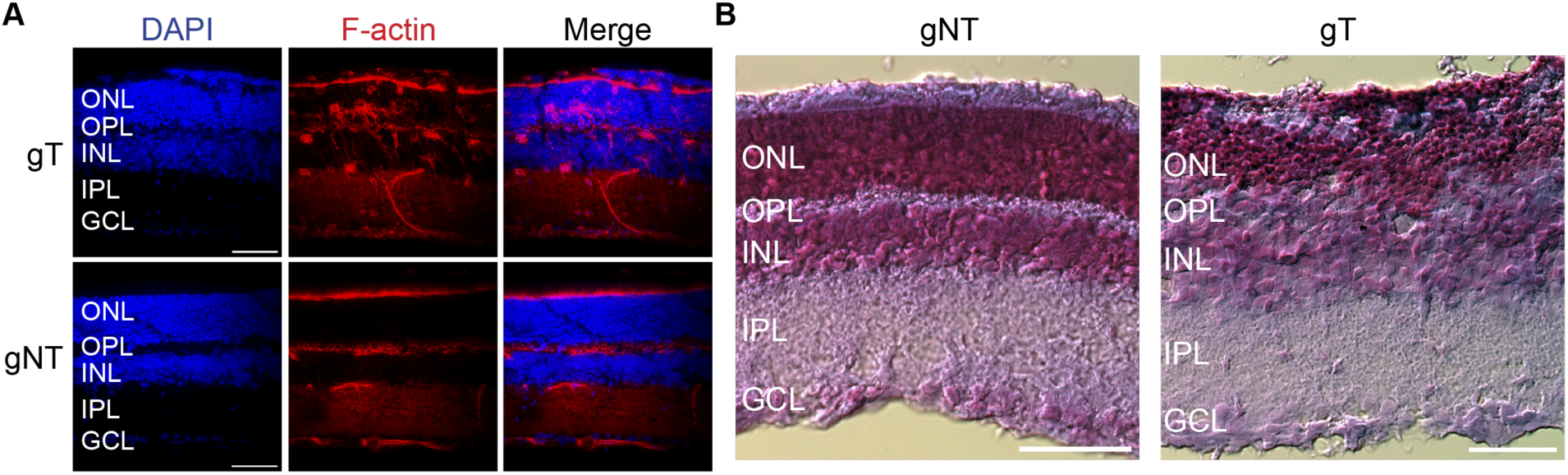
Retinal structural characterization after *Actb* mRNA relocalization. (**A**) F-actin staining for the cytoskeleton. (**B**) H&E staining. Scale bar: 50 µm.

**SI Figure 6.**
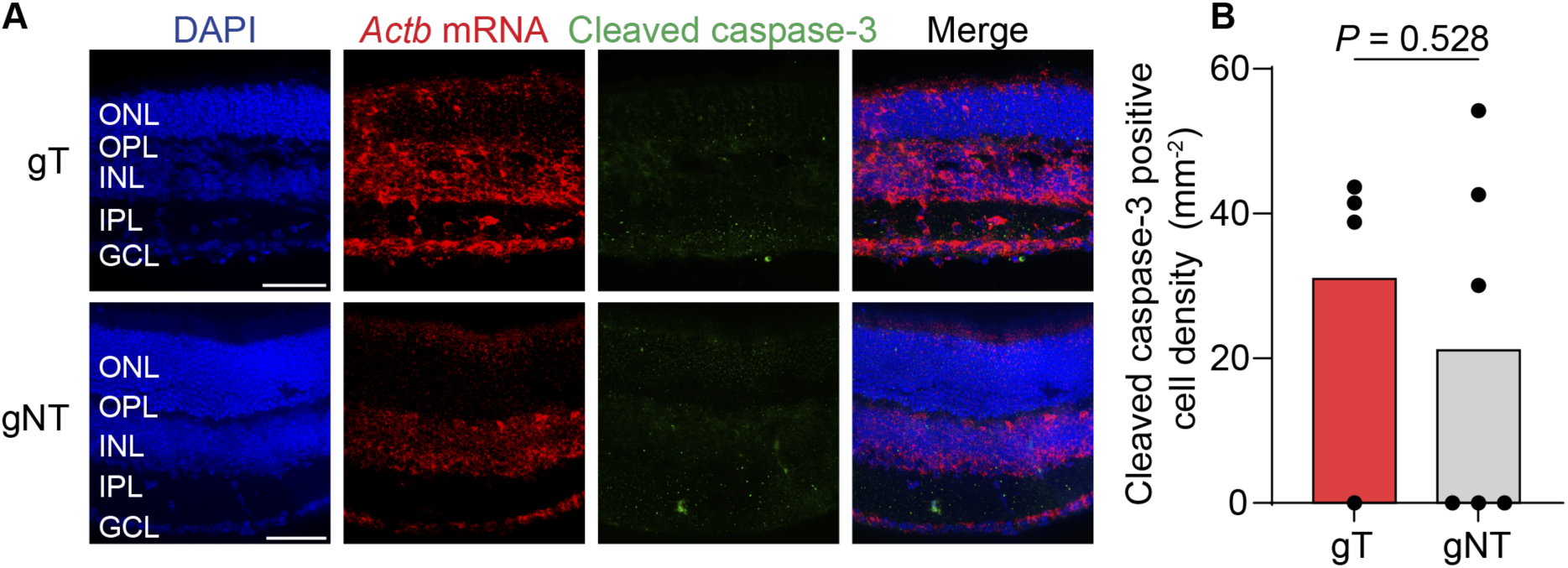
Characterization of apoptosis via cleaved caspase-3 staining after *Actb* mRNA relocalization. (**A**) Representative images of Caspase-3 in retinal slices after 4 h light treatment, overnight dark treatment, and ERG test. The *Actb* mRNA and Caspase-3 for both groups were in the same contrast. Scale bar: 50 µm. (**B**) Statistical result of Caspase-3-positive cell density in the retina. *P* value was calculated using a two-tailed unpaired t-test. From left to right, the number of animals is 4 and 6. Each data point and the mean are shown.

